## Supplementary material for "The distinct role of human PIT in attention control": supplyment

* Sheng He.

**The following includes:**

Figures S1

Figures S2

Tables S1

Tables S2

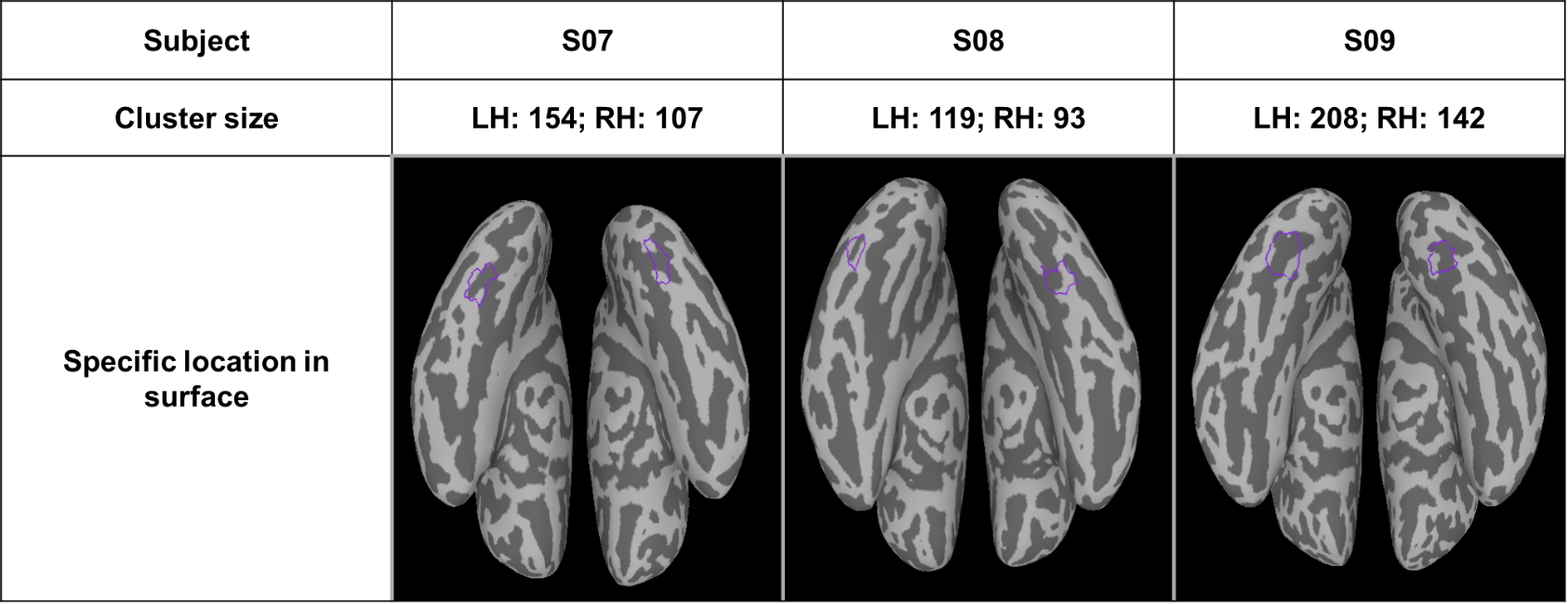

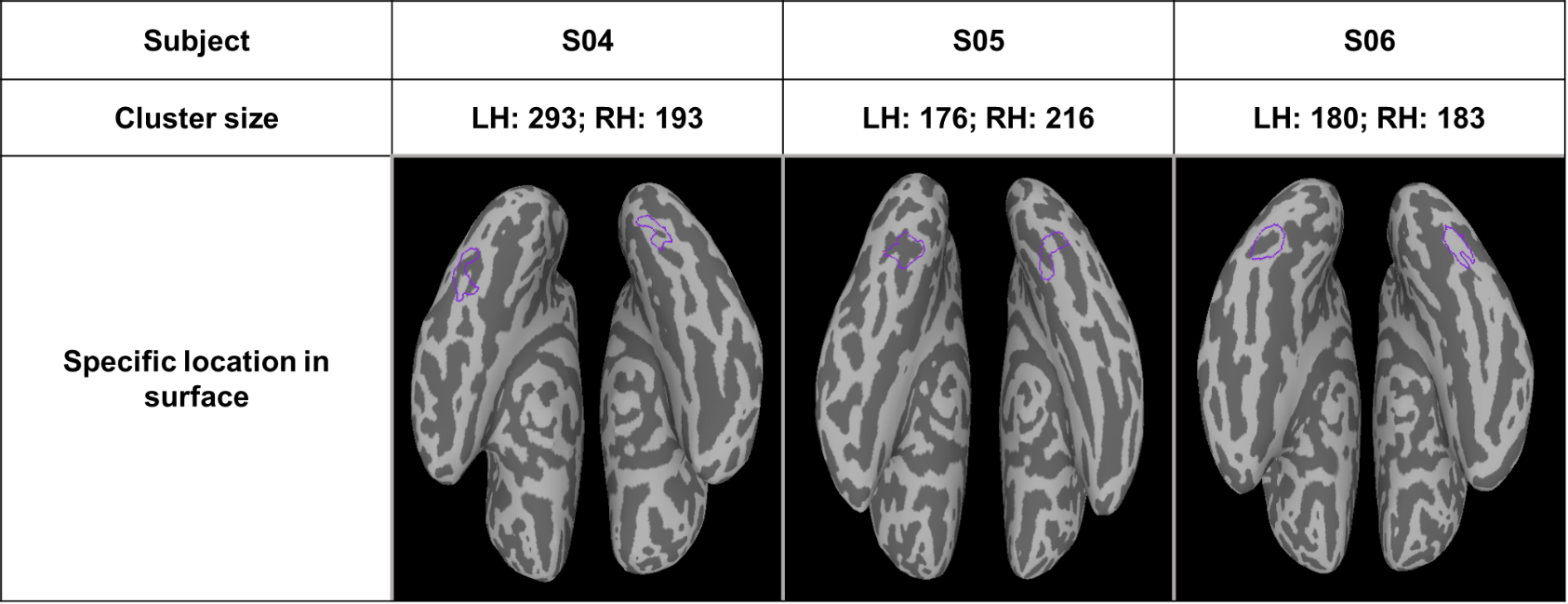

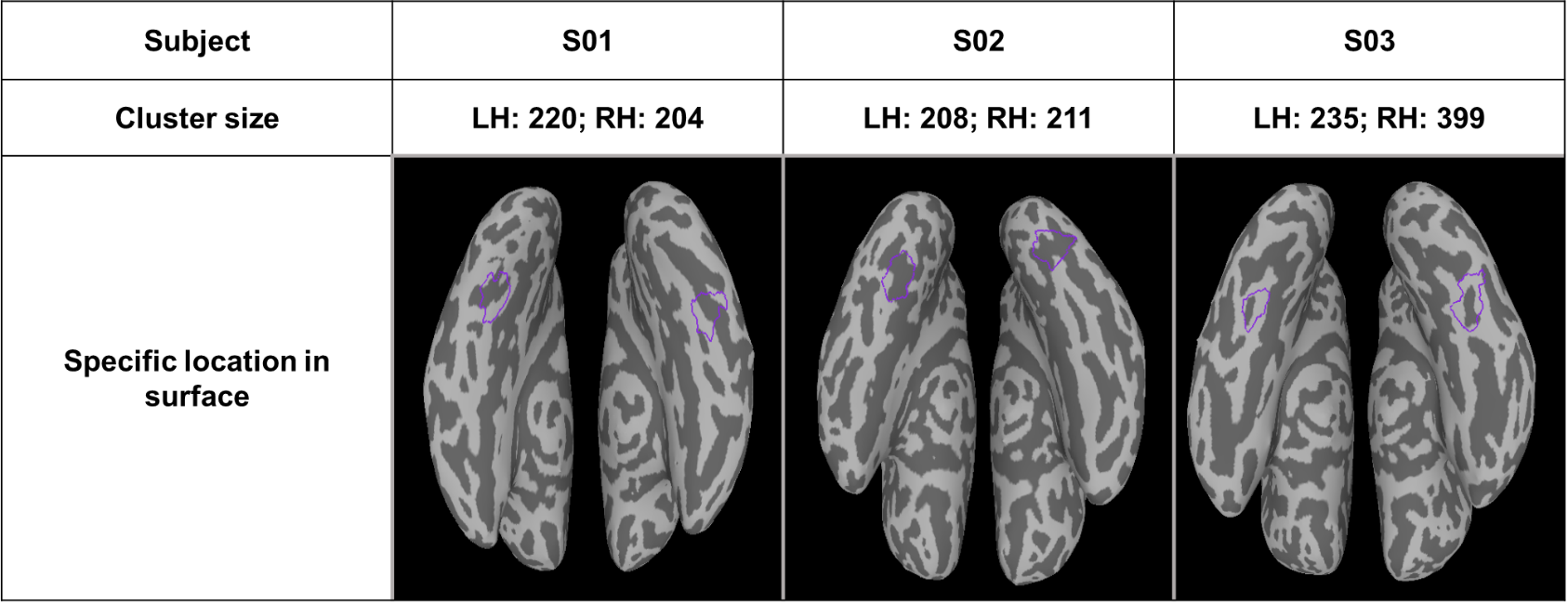

**Fig. S1**
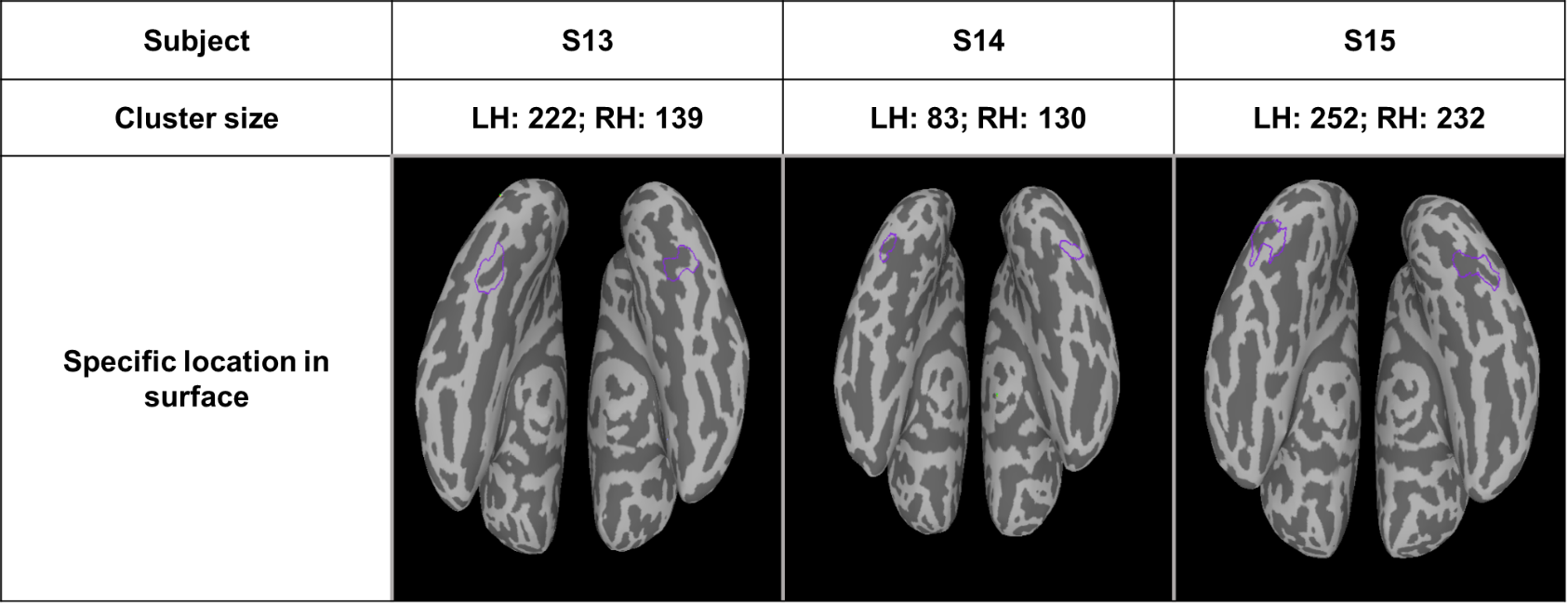

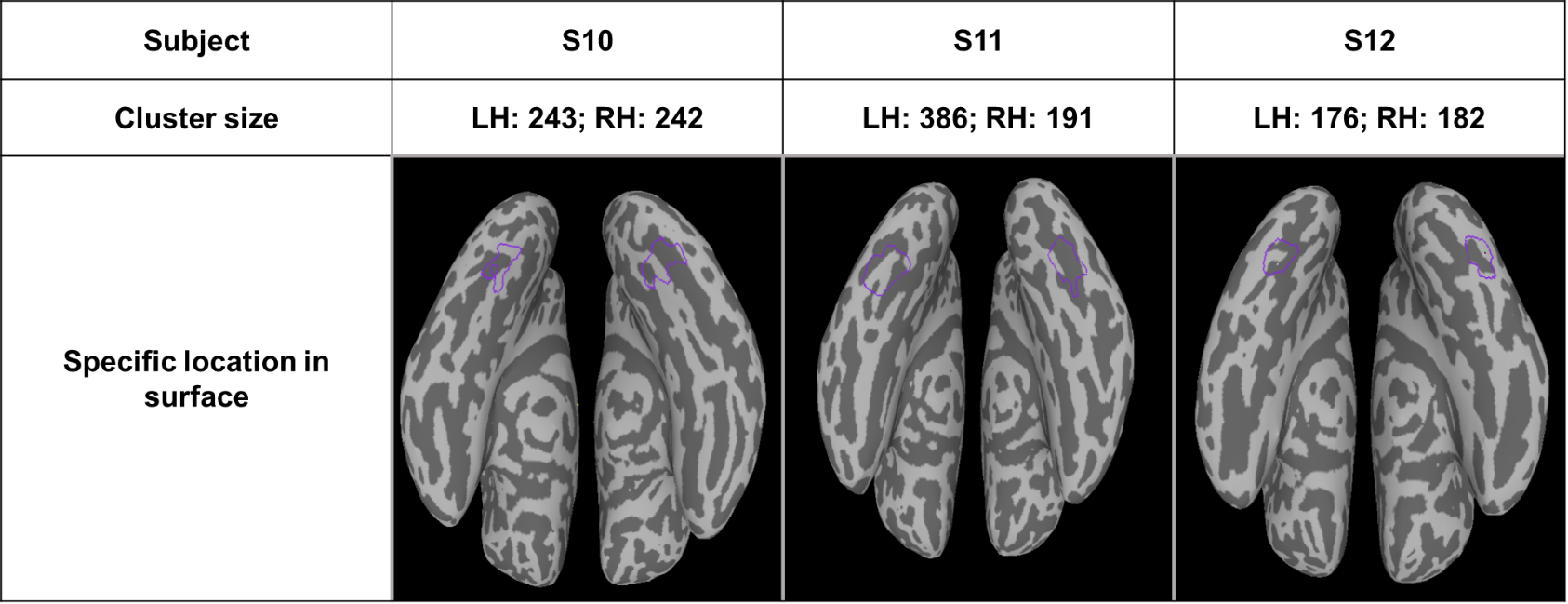
: The exact location and cluster size of hPIT in every subject’s cortical surface. ROI hPIT is selected by purple coil and its cluster size is calculated after projected to volume (one voxel: 2 x 2 x 2 mm).

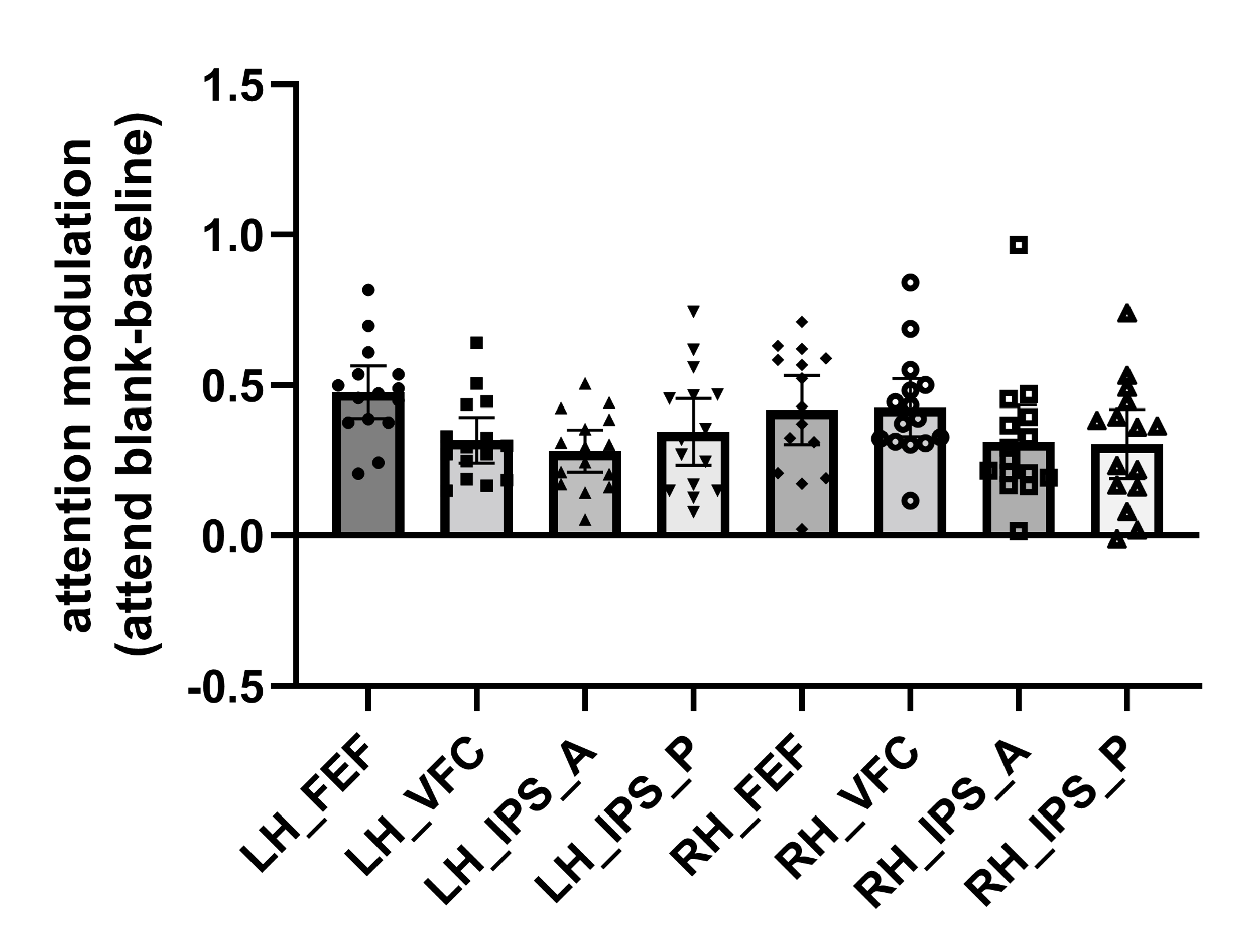

**Fig. S2**: The top-down attentional modulation (attend - baseline) in bilateral frontal and parietal regions during the blank condition. Error bars indicate 95% confidence interval.

| **Condition**  **ROI** | **Blank** | **Dot** |
| --- | --- | --- |
| **hPIT**  **V1**  **MT**  **IPS_P**  **IPS_A**  **FEF**  **TPJ**  **VFC**  **FFA**  **LOp** | 16.59  7.18  9.46  7.57  2.79  3.96  1.89  4.83  3.06  6.75 | 28.44  8.87  16.94  6.73  2.96  3.08  4.10  4.15  19.98  13.36 |

**Table S1**: Summary table of attentional modulation (signal change%) across ROIs (hPIT, V1, MT, IPS, FEF, TPJ, VFC, FFA, LOp) under both blank and dot conditions.

| **Condition**  **ROI** | **Blank**  **(df=14)** | **Dot**  **(df=14)** |
| --- | --- | --- |
| **V1** | t=3.390, p=0.0044 | t=4.743, p=0.0003 |
| **hPIT** | t=4.747, p=0.0003 | t=8.545, p<0.0001 |
| **MT** | t=3.474, p=0.0037 | t=6.162, p<0.0001 |
| **IPS_P**  **IPS_A**  **FEF**  **TPJ**  **VFC** | t=5.975, p<0.0001  t=2.606, p=0.0207  t=5.504, p<0.0001  t=1.756, p=0.1009  t=4.746, p=0.0003 | t=6.035, p<0.0001  t=5.425, p<0.0001  t=5.202, p=0.0001  t=3.196, p=0.0065  t=6.436, p<0.0001 |

**Table S2**: Results of one sample t test measuring the modulation of attention by beta value of contrast [attend contralateral – attend ipsilateral].
